# Interference Length reveals regularity of crossover placement across species

**DOI:** 10.1101/2024.04.22.590575

**Authors:** Marcel Ernst, Raphael Mercier, David Zwicker

## Abstract

Crossover interference is a phenomenon that affects the number and positioning of crossovers in meiosis and thus affects genetic diversity and chromosome segregation. Yet, the underlying mechanism is not fully understood, partly because quantification is difficult. To overcome this challenge, we introduce the interference length *L*_int_ that quantifies changes in crossover patterning due to interference. We show that it faithfully captures known aspects of crossover interference and provides superior statistical power over previous methods. We apply our analysis to empirical data and unveil a similar behavior of *L*_int_ across species, which hints at a common mechanism. A recently proposed coarsening model generally captures these aspects, providing a unified view of crossover interference. Consequently, *L*_int_ facilitates model refinements and general comparisons between alternative models of crossover interference.

## I. INTRODUCTION

Meiotic crossovers (COs) are crucial for ensuring genetic diversity and are necessary for linking maternal and paternal homologs for proper segregation in most eukaryotes. Chromosomes tend to have at least one CO, but rarely more than a handful. Moreover, CO positions are not independent, but exhibit a phenomenon known as *crossover interference* [1–3]: If chromosomes posses multiple COs, they tend to be spaced more widely than expected by chance. The mechanism governing this CO interference is debated [4–16], in part because it is challenging to quantify CO interference reliably and to compare it across species, mutants, and chromosomes.

COs can be detected in cytology using fluorescent imaging of proteins marking CO sites [7, 9, 14, 17–22]; their position then needs to be determined relative to the synaptonemal complex (SC) on which they reside, leading to CO positions quantified in µm in SC space. Alternatively, genetic techniques can detect transmission events from parental DNA to offspring to identify COs [7, 12, 14, 23–27]; positions along the chromatids are quantified in units of megabases (Mb) in DNA space. However, only half of the designated COs will become a CO on a selected gamete [7, 14]. CO maturation inefficiencies can further contribute to discrepancies between the cytologically and genetically obtained data. Both aspects manifest as a random sub-sampling in genetic data [28–30]. Moreover, cytological methods typically detect only class I COs, but not the less prevalent class II COs [7, 11, 13], resulting in a systematic bias.

To quantify CO interference, observed CO counts and positions are summarized using various quantities [13, 23, 28, 36, 37, 40, 41]. In the simplest case, one plots the histogram of observed adjacent distances of COs and compares it to the expected distribution without interference; see Fig. 1A. To obtain a single quantity associated with CO interference, the distribution of distances between adjacent COs can be fitted by a Gamma-distribution; The resulting shape parameter *ν* quantifies the evenness of CO distances and is associated with CO interference [7, 23, 31–38]; see Fig. 1A. However, *ν* is sensitive to random sub-sampling [6, 28, 37, 38], it only uses data from chromosomes with at least two COs [42], and it is also influenced by other aspects than interference, in particular the typically heterogeneous distribution of CO positions [37]. An alternative quantification is the coefficient of coincidence (CoC), which measures the ratio of observed frequency of CO pairs to the expected frequency in absence of interference as a function of the CO distance [2, 3, 14, 38, 39]; see Fig. 1B. The CoC value is close to 1 when interference is absent, but decreases strongly at short distances when interference is present, reflecting the absence of close double-COs. The distance *d*_CoC_ at which the CoC curve crosses 0.5 (orange band in Fig. 1B) provides a length, which tends to be larger for stronger interference [6, 13, 43, 44]. However, this transition point often cannot be located accurately, presumably because it is sensitive to only the data in its vicinity, thus ignoring a potentially large part of the data that could provide information about CO interference. Moreover, the CoC curve relies on binning, which results in information loss [36, 37] and requires the difficult choice of an optimal bin count [12, 28].

**FIG. 1.**
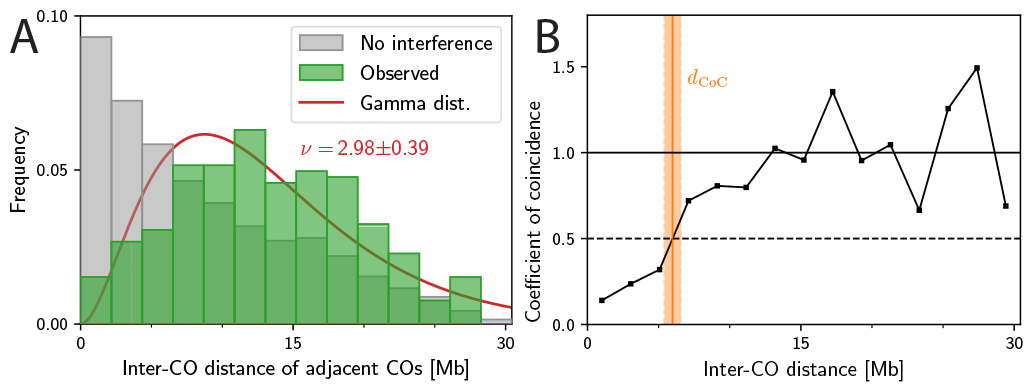
Visualization of traditional quantifications of CO interference. Shown is genetic data from chromosome 1 of wild-type male *A. thaliana* [12]. (A) Comparison of the observed distributions of distances of *adjacent* CO pairs to the expected distribution in absence of interference (obtained by shuffling all CO positions assuming the same distribution of CO count). The indicated shape parameter *ν* follows from a fitted Gamma-distribution (solid line) [7, 23, 31–38]; see SI-A 1. (B) Coefficient of coincidence as a function of the normalized distance between COs [2, 3, 14, 38, 39]; see SI-A 2. The interference distance *d*_CoC_ (orange line, shaded area indicates SEM) marks the point where the curve first exceeds 0.5.

We here introduce the *interference length L*_int_ to complement previous quantifications. After defining *L*_int_ and describing basic properties, we validate it using known behavior of CO interference. We show that *L*_int_ can be used to faithfully compare cytological and genetic data from various species, mutants, and chromosomes. Surprisingly, most of these data can be described by a simple normalized interference length, capturing the regularity of CO positions. This suggests a common mechanism underlying CO interference. Indeed, the recently proposed coarsening model [8, 9, 12, 13] explains this behavior qualitatively.

## II. RESULTS

### A. Interference length: a novel measure for crossover interference

Crossover (CO) interference is quantified based on the observed CO count per chromosome, *N*, and the associated CO positions *x*_*i*_ along each chromosome. One central quantification is the mean number of COs per bivalent, ⟨ *N* ⟩, which is typically reduced when CO interference is strong. However, ⟨ *N* ⟩ does not contain any information about CO positions, so it cannot capture the fact that it is unlikely to find COs in close proximity. To capture such positional information, the main idea of the interference length *L*_int_ is to measures the increase of distances between all (not just adjacent) CO pairs due to CO interference. This increase can be expressed by the difference

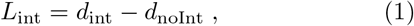

where *d*_int_ quantifies observed distances, with a correction for variations in the distribution of the CO count *N*. In contrast, *d*_noInt_ quantifies the distance in the null hypothesis without interference, where CO are placed independently along the chromosomes, sampling from all observed CO positions. In this null hypothesis, the CO count *N* per chromosome follows a Poisson distribution with the same mean ⟨ *N* ⟩ as the observed data [39]. We define the associated distance *d*_noInt_ as the average distance between any two COs chosen from the pool of all samples for a given chromosome. This definition of *d*_noInt_ preserves the CO density along the chromosome.

To quantify the observed distances and define *d*_int_, we could have simply used the average distance *d*_obs_ of all observed CO pairs. However, this naive choice would only take into account chromosomes with at least two COs, and completely ignore those with one or zero COs. These samples without any CO pairs can represent a large portion of the observation, e.g., in *A. arenosa* [45] and *C. elegans* [5, 7, 46] or in genetic data from *A. thaliana* [12]. In such cases, the naive choice would then only consider data from the small subset with two or more COs, which would dominate the quantification. More importantly, if most samples only carried the obligate CO, strong interference would be likely, which our quantification should capture. Taken together, these arguments show that the distribution of the observed CO count *N* per chromosome carries important information about CO interference.

The observed distribution of CO counts *N* in case of interference is generally narrower than the Poisson distribution of the null hypothesis of no interference; see Fig. 2A. This deviation, even if it is small, can have a significant impact on the number of observed pairs, because there are 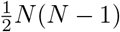 pairs for a chromosome with *N* COs. To see this, imagine observed data of three chromosomes with two COs each, resulting in three distinct pairs; see Fig. 2B. In contrast, without interference, we might have one, two, and three COs on these chromosomes since the distribution of *N* is broader. This would lead to a total of four possible CO pairs, thus providing more pairs than in the observed data, despite identical ⟨ *N* ⟩. This example illustrates that the narrower observed distribution of CO counts *N* leads to fewer CO pairs than the null hypothesis without interference. To account for these *missing pairs*, we compare the average number of observed pairs, 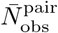, to the average number of pairs in the null hypothesis, 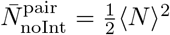, which follows from the assumed Poisson distribution; see SI-B 1. The difference quantifies the average number of missing pairs, 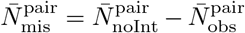. A larger value of 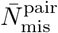 indicates stronger interference, which should be reflected in our measure via a suitable definition of *d*_int_.

**FIG. 2.**
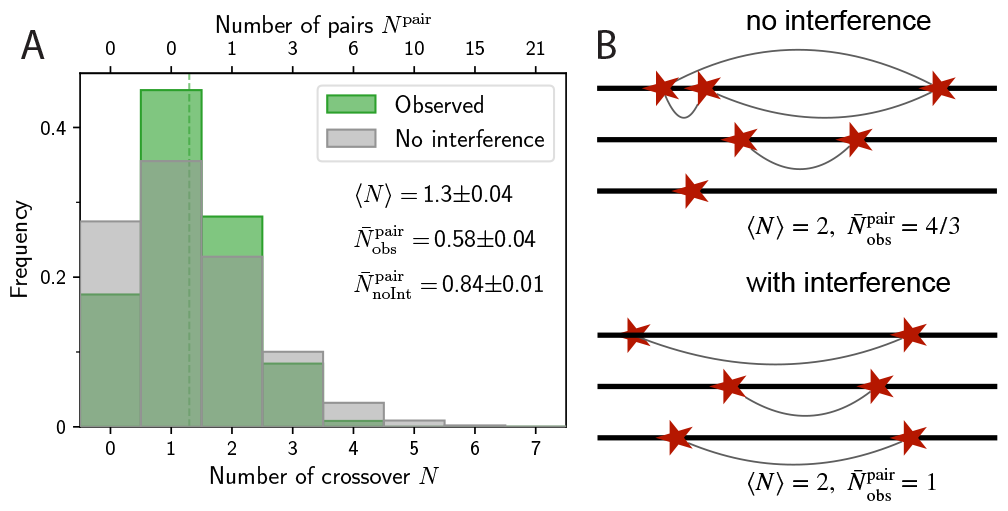
Crossover interference reduces the number of CO pairs. (A) Comparison of the observed distribution (green) of the number *N* of COs per chromosome to the reference without interference (gray) for the same data as in Fig. 1. The corresponding number of CO pairs, 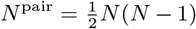, are indicated with respective means. (B) Schematic CO placements on three chromosomes highlighting the effect of interference. The upper panel shows chromosomes with one, two, and three COs, consistent with the broad Poisson distribution in the case without interference. In contrast, interference typically leads to a narrower distribution (bottom panel), where each chromosome has two COs. While both cases have the same mean CO count, ⟨ *N* ⟩= 2, the thin gray lines indicate that we have a total of three CO pairs with interference 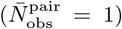, and thus less than in absence of interference 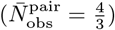, suggesting interference reduces 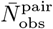.

The distance *d*_int_ quantifies the distance of CO pairs in case of interference, which should capture the actually observed distances as well as the fact that interference is stronger when there are more missing CO pairs. We thus define *d*_int_ using a weighted average of observed and missing pairs,

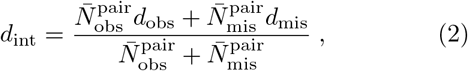

where *d*_obs_ is the mean distance between all (not just adjacent) CO pairs on the same chromosome. In contrast, *d*_mis_ quantifies the distance associated with missing pairs, and we here choose the chromosome length *L* as the most natural length scale, *d*_mis_ = *L*. We will discuss below how this choice is related to the maximal interference length that can realistically be observed. Taken together, the interference length can be expressed as

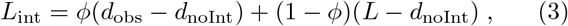

Where 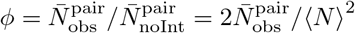 denotes the ratio of observed to expected CO pairs, which is small in case of strong interference; compare Fig. 3A. Eq. (3) highlights that the interference length *L*_int_ combines information of (i) the distribution of CO positions via *d*_noInt_, (ii) the distribution of the observed distances of CO pairs via *d*_obs_, and (iii) the distribution of observed CO counts via *ϕ*.

**FIG. 3.**
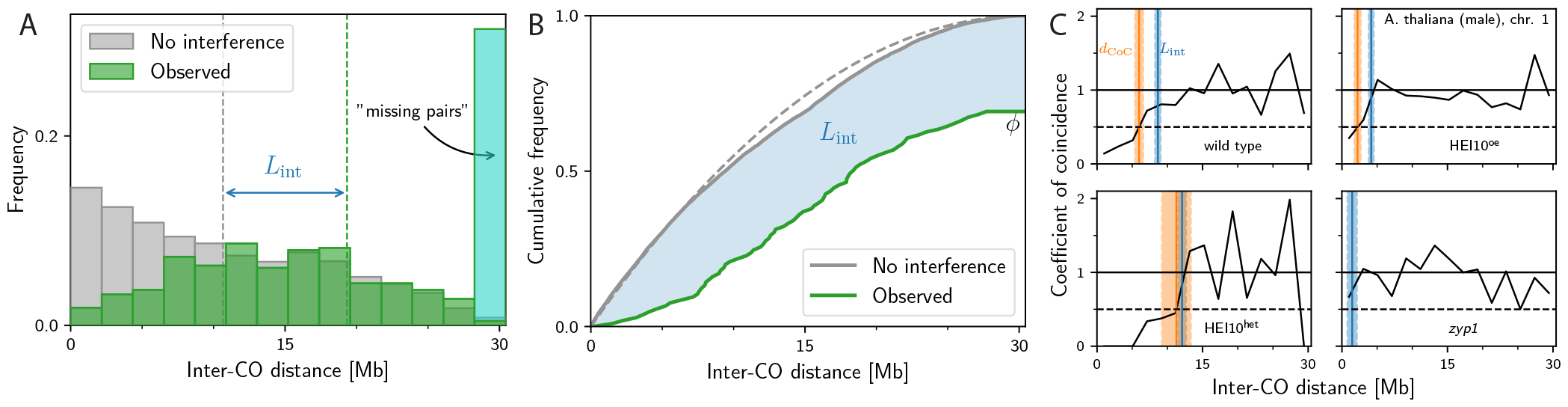
Visualizations of interference length *L*_int_. Shown is data for the first chromosome of male meiosis of *A. thaliana* [12]. (A) Comparison of the observed (green) and expected (gray) distribution of distances of all CO pairs for wildtype data. The last, cyan bin accounts for missing pairs, which contribute with a length of *d*_mis_ = *L* where *L* = 30.4 Mb is the measured chromosome length. The interference length *L*_int_ is the distance between the mean values of these distributions (denoted by vertical dashed lines). (B) Cumulative distributions corresponding to panel A. The blue area between the curves corresponds to *L*_int_. The dashed gray line indicates the theoretical distribution for uniform CO distributions. (C) Coefficient of coincidence curves of four different genotypes of chromosome 1 of male *A. thaliana* [12]. Vertical bands mark associated interference lengths *L*_int_ (blue) and interference distances *d*_CoC_ (orange) with respective SEM.

Fig. 3A shows a graphical interpretation of the interference length *L*_int_ based on the histogram of the distances between all CO pairs per sample. In contrast to Fig. 1A, we account for missing CO pairs, which contribute with the maximal distance *L* (cyan region). Consequently, the mean distance of the observed data shifts to larger values (compare dashed green lines in Fig. 1A and Fig. 3A), capturing that missing CO pairs indicate strong interference. Fig. 3B visualizes the same idea using cumulative distribution functions. Here, *L*_int_ corresponds to the blue area between the gray curve representing the null hypothesis and the green curve for observed data with interference, which is scaled by *ϕ* to account for missing CO pairs. The cumulative distribution function highlights that *L*_int_ can be determined without binning, abolishing this step that could degrade data quality.

The interference length *L*_int_ has multiple properties that make it a suitable measure of CO interference: (i) *L*_int_ is a scalar quantity of dimension *length*. Consequently, *L*_int_ is reported in units of µm for cytological data (SC space), and units of megabases (Mb) for genetic data (DNA space). (ii) We show in SI-B 2 that *L*_int_ is invariant to random sub-sampling (similar to CoC curves), which facilitates the comparison of cytological and genetic data. (iii) *L*_int_ uses all empirical data on CO positions and does not use any binning or parametrization. On the one hand, all observed CO pairs contribute equally to *d*_obs_ and thus *L*_int_; see Eqs. (2)–(3). On the other hand, the definition also accounts for chromosomes without COs or only one CO via the average number of missing pairs,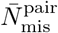. (iv) The quantity *d*_noInt_ is based on the observed distribution of CO positions along the chromosome, so that variations of CO density, e.g., due to suppression in centromeric regions, are incorporated in *L*_int_. (v) *L*_int_ allows for uncertainty estimations (SI-B 3) and significance testing (SI-B 4). We provide a reference implementation of *L*_int_ with the Supporting Material.

### B. Large interference lengths indicate strong interference

To see how well the interference length *L*_int_ captures CO interference, we start by comparing it to the more traditional CoC curves. Fig. 3C shows four representative CoC curves for various strains of *A. thaliana*, known to exhibit very different CO interference. In all cases, *L*_int_ (blue bands) qualitatively captures the distance at which the CoC curve approaches 1, indicating the point at which distances between COs are as frequent as in the null hypothesis without interference. In particular, *L*_int_ is larger for cases known to exhibit strong interference (e.g., the HEI10^het^ mutant), and it correlates (cf. Fig. 9C) with the interference distance *d*_CoC_ (orange band; where the CoC curves exceeds 0.5). However, *L*_int_ can be calculated more precisely (indicated by the smaller standard error of the mean; see Fig. 7B), and it can also be determined for cases without interference (e.g., the *zyp1* mutant) and when few CO pairs are observed. This first analysis thus indicates that *L*_int_ captures essential aspects of CoC curves and CO interference.

The only crucial parameter in the definition of *L*_int_ is the distance *d*_mis_ associated with missing pairs. Our choice of *d*_mis_ = *L* implies that *L*_int_ assumes values on the order of the chromosome length *L* in cases of strong interference. Since there are multiple cases that could be called “strong interference”, we next evaluate *L*_int_ for four theoretical scenarios: (i) When all chromosomes exhibit exactly one CO per chromosome, we have *ϕ* = 0 and thus *L*_int_ = *L− d*_noInt_. In this scenario of *complete interference*, we obtain 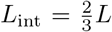 when COs are distributed uniformly along the chromosome; see SI-B 5. These results persist if some chromosomes have no CO instead of one. (ii) We also find 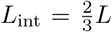 when all chromosomes have exactly two COs at opposite ends of the chromosome. (iii) The *maximal-interference model* of *L*_int_ for a given average CO count *N* yields 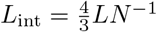 (limited to *L*_int_ = *L* for *N* = 1 when COs always occur at the same position); see SI-B 6. (iv) Finally, we consider the case where exactly *N* COs are placed at fixed distance *L/N*, and the first CO is located uniformly between 0 and *L/N*, so the overall CO frequency is uniform along the chromosome. This *regular-placement model* predicts 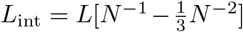; see SI-B 7. Taken together, these theoretical scenarios suggest two limiting behaviors of *L*_int_ in case of strong interference: For few COs, ⟨ *N* ⟩ ≈ 1, the first two scenarios suggest 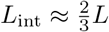. Conversely, for many COs, ⟨ *N* ⟩ ≫1, the last two scenarios suggest the scaling *L*_int_∼*L/* ⟨ *N* ⟩ We expect that intermediate values of *N* interpolate between these two extremes. In the contrasting case without interference, when the CO count *N* follows a Poisson distribution and COs are placed independently (but not necessarily uniformly) along the chromosome, we have *ϕ* = 1, *d*_obs_ = *d*_noInt_, and hence *L*_int_ = 0, corresponding to the null hypothesis without interference. This indicates that larger values of *L*_int_ are associated with stronger interference, and that the precise value depends on *L* and ⟨ *N* ⟩. We next test these predictions for experimental data.

### C. Interference length recovers sex differences and mutant behavior

We start by using the interference length *L*_int_ to query known properties of CO interference across different chromosomes, genotypes, and species. One advantage of *L*_int_ is that cytological and genetic data can be compared directly because *L*_int_ is invariant to sub-sampling. To test this property explicitly, we took advantage of published data where both genetic and cytological data where available. This includes human male [49, 50], as well as *A. thaliana* wild type and mutants with variations in the expression levels of HEI10 [9, 12, 47, 48]; see details of data handling in SI-C. Fig. 4A shows that the average CO count ⟨ *N* ⟩ of the cytological data is approximately twice that of the genetic data. This is consistent with expected sub-sampling since a CO detected in cytology affects only two of the four chromatids, and is thus detected in only half the gametes [28–30].

**FIG. 4.**
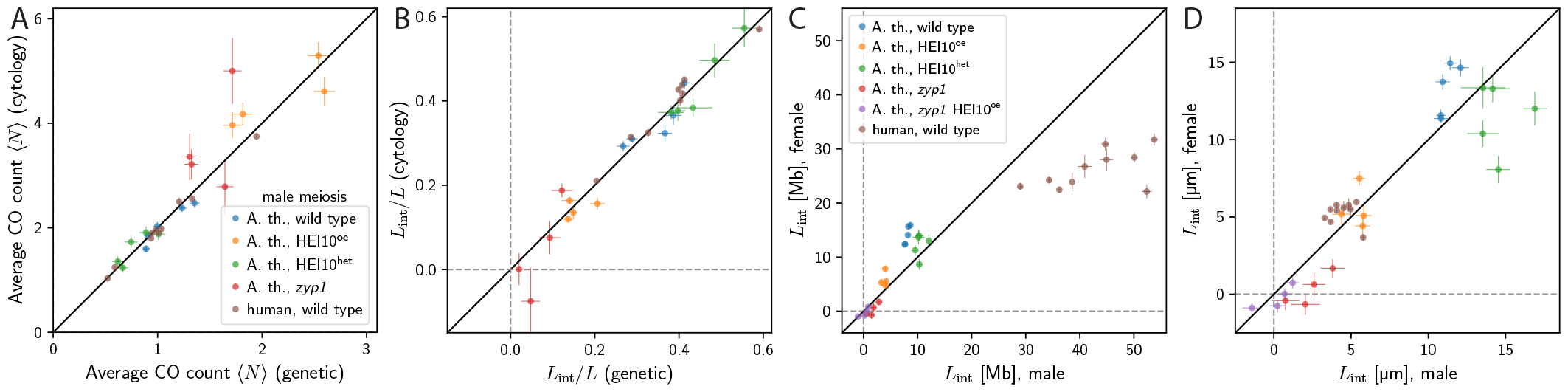
Interference length retrieves known results. (A) Comparison of the average CO count ⟨ *N* ⟩ for male meiosis in *A. thaliana* for various genotypes based on genetic [12, 47] and cytological data [9, 48], and male human wild type based on genetic [49] and cytological data [50] for individual chromosomes. The black line indicates the expectation that ⟨ *N* ⟩ is twice as large for cytology compared to genetic data. (B) Comparison of the interference length *L*_int_ normalized to the chromosome length *L* for the same data as in (A). (C) Comparison of *L*_int_ for male and female meiosis for various genotypes based on genetic data of *A. thaliana* [12, 47] and wild-type, as well as cytological data for human [50] scaled with the respective DNA lengths according to [51] and thus measured in DNA space [Mb]. (D) Comparison of *L*_int_ of the same data as in (C); *A. thaliana* data is scaled with respective SC lengths [12, 50] and thus measured in SC space [µm]. (A–D) Details of data handling in SI-C.

We next compare the interference lengths *L*_int_*/L* determined for the genetic and cytological data normalized with the chromosome length and the SC length, respectively. Fig. 4B shows that cytological and genetic data lead to very similar values of *L*_int_*/L*. In particular, the null hypothesis that the values agree is not rejected for *A. thaliana* wild type (*p* = 0.95, significance test described in SI-B 4), HEI10^oe^ (*p* = 0.50), HEI10^het^ (*p* = 0.84), and *zyp1* (*p* = 0.69), as well as human (*p* = 0.10).

Another important feature of CO interference are sex differences, where CO rates differ between female and male. In *A. thaliana*, female meiosis generally features fewer COs and stronger CO interference according to coefficient of coincidence (CoC) analysis [9, 12, 21, 27]. Fig. 4C shows that genetic data [12, 47] of females indeed exhibit larger interference lengths *L*_int_ in DNA space than males. This difference is significant in wild type (*p* = 10^−4^), but not in HEI10^oe^ (*p* = 0.06) and in HEI10^het^ (*p* = 0.32). It is generally accepted that interference propagates in the µm space of the SC [6, 21]. Indeed, when we convert the genetic data from DNA space to SC space using the chromosome and SC lengths reported in ref. [12] and then calculate *L*_int_, the difference between female and male is less significant for *A. thaliana* wild type (*p* = 0.02) and is absent for HEI10^het^ (*p* = 0.09) as well as HEI10^oe^ (*p* = 0.83) see Fig. 4D. Taken together, this supports a common process in male and female governing CO interference in SC space, whereas sex differences are a consequence of different chromosome organisation, consistent with literature [6, 21].

To corroborate this, we also investigated sex differences for human data from cytological imaging of MLH1 foci [50], where CoC analysis in SC space suggest no significant sex difference, thus implying weaker interference for females if measured in DNA space due to lower DNA compaction in female meiosis [50]. Instead, we find a weakly significant difference for *L*_int_ for the cytological data (Fig. 4D, *p* = 0.03), whereas converting cytological data from SC space to DNA space using chromosome lengths reported in ref. [51] results in significantly smaller *L*_int_ for females (Fig. 4C, *p* = 10^−6^). Our analysis again suggests that sex differences are predominately caused by different chromosome compaction, whereas female and male exhibit similar CO interference in SC space.

Finally, we test whether *L*_int_ recovers the behavior of *A. thaliana* mutants. Increasing HEI10 levels (HEI10^oe^ line) decreases *L*_int_ for both male (*p* = 10^−3^) as well as female (*p* = 4 ·10^−4^) in genetic data [12] and for male cytological data (*p* = 2 ·10^−4^) [45]; see Fig. 4B– D. Lowering HEI10 levels (HEI10^het^ line) increases *L*_int_ for male genetic data (*p* = 0.04) [12] and cytological data (*p* = 0.045) [45], but *L*_int_ remains unchanged for female genetic data (*p* = 0.15), suggesting that CO interference is already almost maximal (*L*_int_*/L* = 0.52 … 0.67). For mutants where the SC is absent [12, 27, 48, 52], *L*_int_ is consistent with absent interference in female *zyp1* mutant [12] (*p* = 0.60), the male *zyp1* mutant (cytology) [48] (*p* = 0.23) and the double mutant *zyp1* HEI10^oe^ [12] (male *p* = 0.56, female *p* = 0.78), whereas male *zyp1* mutants (genetic) [12] might exhibit some residual interference (absent with *p* = 0.04). We thus showed that the interference length *L*_int_ recovers known behavior of CO interference in *A. thaliana* mutants.

### D. Interference length facilitates comparison across multiple species

We established that the interference length *L*_int_ tends to be larger when CO interference is stronger and that this correlation recovers many aspects of CO interference. However, we so far have not interpreted the numeral value of *L*_int_ in detail, particularly when comparing different genotypes or even different species. Since *L*_int_ is a single number, such a comparison is easily feasible and can shed light onto the mechanism of CO interference in different species.

To compare measured interference lengths *L*_int_ of different species, we show *L*_int_ obtained from cytological data as a function of the SC length *L* in Fig. 5A. Evidently, *L*_int_ can vary widely across species, even when SC lengths are comparable. For instance, for *L*·≈ 40 µm, *A. arenosa* exhibits *L*_int_ ≈20 µm, whereas *A. thaliana*, maize, and human exhibit progressively smaller values down to *L*_int_ ≈ 5 µm, suggesting reduced CO interference. However, we also find that *L*_int_ is correlated with *L*: Multiple species (*A. arenosa* [45], *C. elegans* [46], mouse [54], and tomato [18]) exhibit data very close to the line 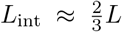, which we associate with complete interference motivated by the theoretical scenarios studied above. Whereas these species exhibit an almost proportional relationship between *L*_int_ and *L*, other species (maize [53], *A. thaliana* [9], and human [50]) exhibit a weaker dependence. The associated values of *L*_int_ are smaller than 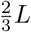, indicating incomplete interference. However, all observed wild-type values are significantly larger than zero, suggesting that they all exhibit CO interference. Taken together, this initial comparison suggests that species either exhibit strong interference close to maximal values 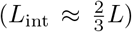 or they exhibit smaller values and weaker *L*-dependence.

We next investigate how mutations change *L*_int_ for a few species. Fig. 5B. shows that the *C. elegans ie29* strain (green triangle) has the same value of *L*_int_ as the wild type (*p* = 0.68), suggesting that this strain does not exhibit altered CO interference. In contrast, *L*_int_ is strongly reduced for *C. elegans syp-4* mutants (green circle [46]), consistent with the idea that an intact SC is required for CO interference. We observe a similarly strong reduction of *L*_int_ in *A. thaliana zyp1* mutants (orange circles), consistent with the described abolished interference [9, 48]. In *A. thaliana, L*_int_ can also be reduced by over-expressing HEI10 (orange triangles pointing up), whereas interference is increased when HEI10 levels are reduced in the HEI10^het^ strain (orange triangles pointing down), consistent with the analysis of the genetic data shown above and literature [9, 12, 48].

**FIG. 5.**
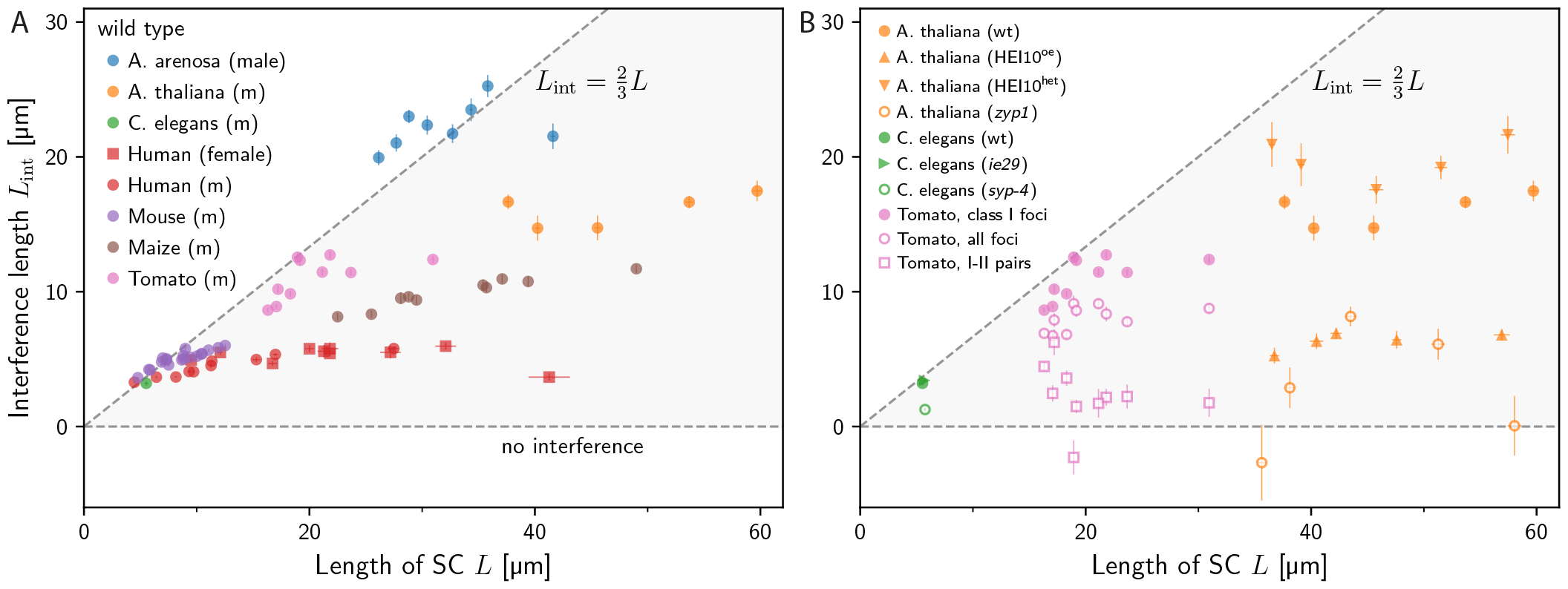
Interference length allows for simple comparison across species and genotypes. (A) Interference length *L*_int_ as a function of SC length *L* for cytological data of wild-type data of *A. arenosa* [45], *A. thaliana* [9], *C. elegans* [46], human [50], maize [53], mouse [54], and tomato [18]. (B) Interference length *L*_int_ as a function of SC length *L* for cytological data of indicated genotypes for *A. thaliana* [9, 48], *C. elegans* [46], and tomato [18]. For tomato, we present the interference length of class I COs, of all observed foci (class I and class II CO), as well as pairs with one class I and one class II CO (cf. SI-B 9). (A–B) Details of data handling in SI-C.

A challenge in interpreting CO interference experimentally is that some methods (e.g., based on labeling MLH1 in cytology) only observe class I COs, whereas others (e.g., based on electron microscopy or genetics) cannot distinguish class I COs from class II COs [7, 11, 13]. A study in tomato [18] used correlative microscopy to identify MLH1-positive recombination nodules (class I CO) and MLH1-negative nodules (class II CO) in the same cells. We analyzed these data and determined *L*_int_ for various combinations of the two classes of COs; see Fig. 5B. The resulting *L*_int_ is largest when it is determined only for class I COs (pink disks), which are known to exhibit interference. The value reduces significantly (*p*≈10^−4^) when *L*_int_ is calculated based on all foci (pink circles), and this reduction is consistent (*p* = 0.29) with an approximate correction of *L*_int_ taking class II COs (6% to 19%) into account; see SI-B 8. We also quantify how class II COs interfere with the positioning of class I COs by evaluating *L*_int_ associated with pairs comprising a class I CO and a class II CO (pink squares); see SI-B 9. These mixed pairs exhibit a positive (*p* = 0.01), weaker interference (*p* = 10^−4^, compared with *L*_int_ of all foci), but we have no evidence that class II COs interfere with each other (*p* = 0.58, *L*_int_ not shown in figure) or exhibit different interference than the mixed pairs (*p* = 0.51), consistent with literature [18].

Taken together, these data show that the interference length *L*_int_ recovers central observations about CO interference. In particular, values of *L*_int_ tend to vary between small values (indicating absence of interference) and large values 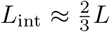 (indicating strong interference). While we briefly explored the dependence on the chromosome length *L*, we expect from our theoretical analysis that *L*_int_ also depends on the mean CO count ⟨ *N* ⟩, which could distort the interpretations we made so far.

### E. Crossover interference exhibits similarity across species and mutants with intact SC

The maximal-interference model and the regular-placement model suggest the scaling *L*_int_∼*L/* ⟨ *N* ⟩, i.e., that *L*_int_ is generally larger for longer chromosomes and fewer COs. To test this scaling, we analyze the normalized interference length, 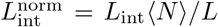, which would be a constant if the scaling held perfectly. Since *L/* ⟨ *N* ⟩ estimates the expected distance between COs, 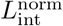 relates to the regularity of COs placement along chromosomes. Note that ⟨ *N* ⟩ is the number of COs per bivalent, implying that we need to double the CO counts measured for individual chromatids in genetic data to account for the sub-sampling. Fig. 6A shows that 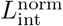 clusters around values between ∼0.6 and ∼0.8 for wild types of many species, particularly when they have few COs (⟨ *N* ⟩ ≲ 4). A notable exception is *S. cerevisiae*, which generally seems to exhibit weaker CO interference than other species we analyzed.

**FIG. 6.**
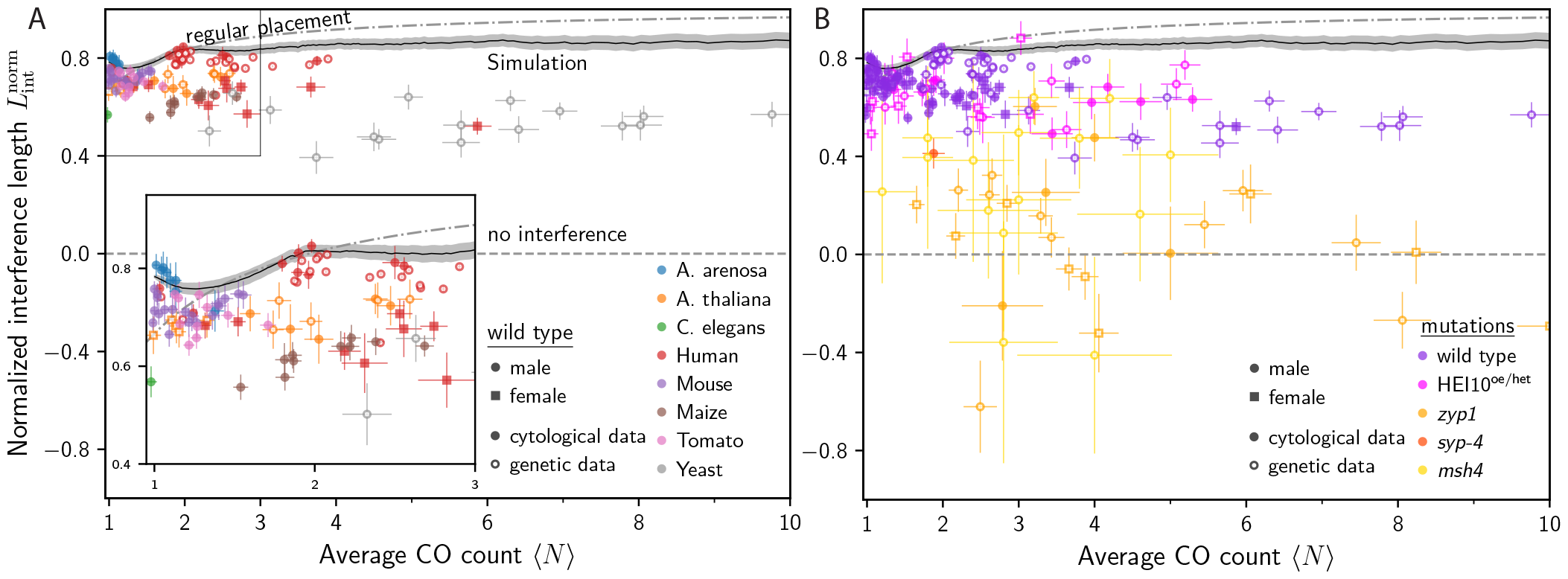
Normalized interference length unveils similarity of mutant behavior across species. (A) Normalized interference length 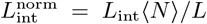 as a function of the mean CO count ⟨*N* ⟩ for wild-type data of *A. arenosa* [45], *A. thaliana* [9, 12], *C. elegans* [46], human [49, 50], maize [53], mouse [54], tomato [18], and *S. cerevisiae* [55] using both cytological and genetic data. (B) 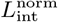 as a function of *N* for the same wild-type data as in Panel A (violet), mutations with altered HEI10 levels in *A. thaliana* (magenta, [9, 12, 47]), mutations that affect the SC in *A. thaliana* and *C. elegans* (orange, [12, 46, 48]) and the *msh4* mutant for budding yeast (gold, [55]). (A-B) The dashed line marks the prediction of the regular-placement model corresponding to strong interference (see SI-B 7), whereas the black line corresponds to the coarsening model for *A. thaliana* [12]. Details of data handling in SI-C.

To explain the observed narrow band of 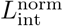, we compare the data to two theoretical predictions. First, we investigate the regular-placement model (dashed lines in Fig. 6), where ⟨ *N* ⟩ COs are placed uniformly with separation *L/*⟨ *N* ⟩ This model overestimates *L*_int_ for larger values of ⟨ *N* ⟩, likely because CO placement is not as regular in reality. Interestingly, the model underestimates *L*_int_ for small ⟨ *N* ⟩, which is a consequence of its uniform CO placement along the chromosome, whereas the observed distributions are often highly non-uniform. Second, we study the recently proposed coarsening model of CO interference [8, 9, 12] for parameters obtained for *A. thaliana* [12]. While the model (black lines in Fig. 6) captures the general trend better than the regular-placement model, there are significant deviations: The model overestimates 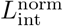 for most species, except *A. arenosa* [45], most likely because of very localized CO positions. The discrepancies between data and model revealed by 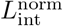 could guide future model refinements.

Our analysis of the normalized interference length 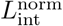 for simple models and wild-type data suggests that systems with strong interference exhibit similar values of 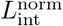, which depend only weakly on *L* and ⟨ *N* ⟩ In particular, the normalized interference length removes the dependency on *L* and ⟨*N* ⟩ that dominated in Fig. 5A: On the one hand, the species obeying the scaling 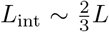 all exhibit ⟨ *N* ⟩ ≈ 1, implying that the associated 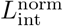 is roughly 0.7. On the other hand, the cases in Fig. 5A that deviated from this scaling all exhibit more COs, explaining the reduced values of *L*_int_. Consequently, all the cases shown in Fig. 5A (except human female chromosome 1 with ⟨ *N* ⟩ ≈ 6) exhibit values of 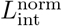 between ∼0.6 and ∼0.8. This similarity in 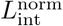 in all analyzed species (except budding yeast, which exhibits larger CO counts and lower *L*_int_) indicates a similar regularity in CO placement, which could originate from a similar mechanism that governs CO interference in these different species.

We next test the hypothesis that the normalized interference length 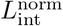 captures an essential aspect of the CO interference process by comparing wild-type data with mutants known to affect CO interference. Fig. 6B shows that mutants affecting the SC (orange and gold symbols) exhibit lower values of 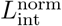 which seem to cluster around 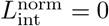 This observation is consistent with the strongly reduced interference described in the literature [12, 46, 48, 55], which disrupts the regularity of CO placement. In contrast, *A. thaliana* mutants with altered HEI10 levels (magenta symbols) exhibit values of 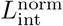 that are consistent with the wild-type results (violet symbols). Apparently, changing HEI10 levels only affects the CO count *N* but not the CO interference as measured by 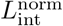 Taken together, we propose that 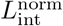 quantifies aspects of CO interference that are independent of ⟨ *N* ⟩, which suggests that CO interference is not affected by changing HEI10 levels, but is strongly impaired in mutants affecting the SC. This interpretation is consistent with the coarsening model, where the SC is vital for mediating coarsening between COs on the same chromosome, whereas changing HEI10 levels merely affects the degree of coarsening without disrupting the mechanism.

## III. DISCUSSION

In this paper we propose the *interference length L*_int_, which summarizes deviations in CO placement, as a new quantity to measure CO interference. *L*_int_ is a physical length, which is larger for stronger CO interference and reaches a maximum of about 0.8*L* in empirical data. The fact that *L*_int_ provides a single number to measure CO interference enables direct comparison of data for different chromosomes, genotypes, and species among each other and with theoretical models. The quantity is also invariant to random sub-sampling (enabling comparison of genetic and cytological data), does not require binning, and uses all empirical data, particularly those from chromosomes with one or no COs. The uncertainty of *L*_int_ is lower than comparable quantities (like *d*_CoC_ or *ν*, see Fig. 7B), providing more statistical power at the same sample size or allowing for fewer experiments to draw conclusions.

**FIG. 7.**
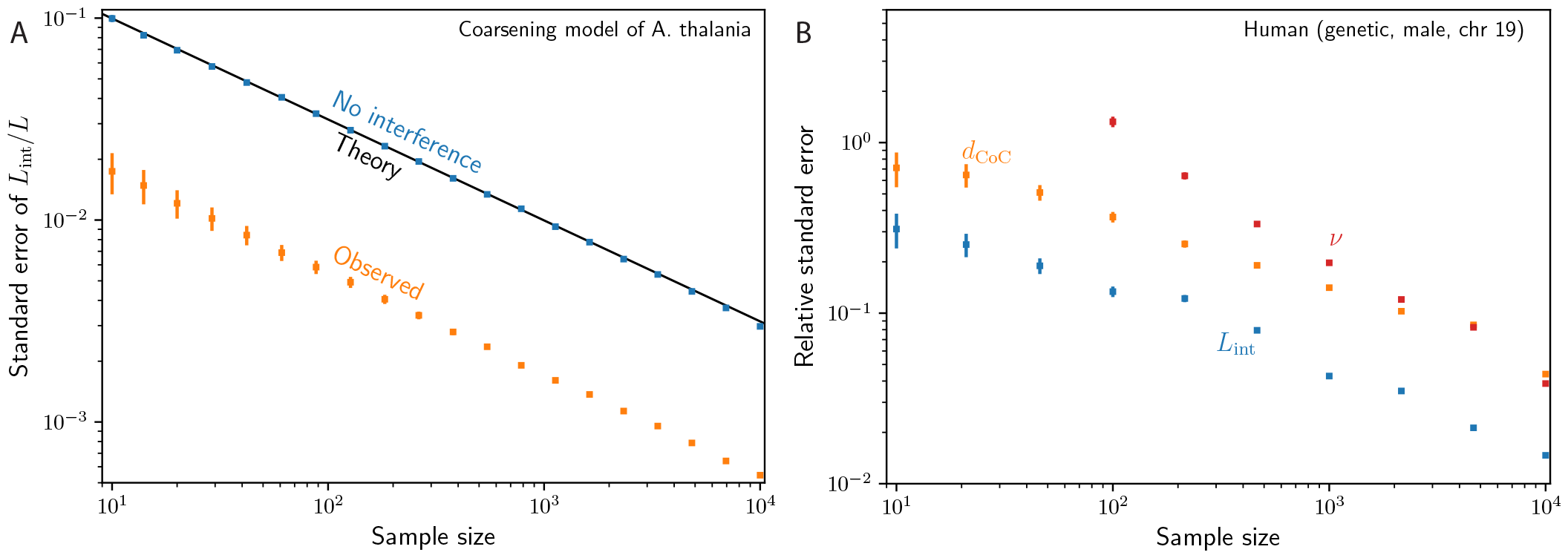
Uncertainty of interference length *L*_int_ is smaller compared with traditional quantifications. (A) Uncertainty (standard deviation of the mean) based on bootstrap sampling for the normalized interference length (*L*_int_*/L*) as a function of sample size in (i) the case of absent interference by simply assuming uniform distribution of crossovers along the chromosome (grey) and (ii) the case of crossover interference by applying the coarsening model in [12] (50000 overall samples) for a chromosome of length 50 µm (green). In both cases we have an average of *N*≈ 3.2 crossovers per chromosome. We add a fit with a slope of −1*/*2 for the case of crossover interference (grey) and the theoretical uncertainty for absent interference (black). (B) Uncertainty based on bootstrap sampling for human genetic data (31228 overall samples) for chromosome 19 which has an average CO count of ⟨ *N* ⟩ ≈ 1. Shown is the standard deviation as a function of sample size of the (i) normalized interference length *L*_int_*/L* (blue) (ii) normalized interference distance *d*_CoC_*/L* (orange) using 15 intervals to compute the CoC curves (iii) gamma shape parameter *ν* normalized with the actual *ν* based on the full data set.

We used *L*_int_ to query known behavior of CO interference using published data. The comparison across species revealed that *L*_int_ only reaches maximal values when there are few COs 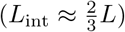 In contrast, species with larger CO counts typically exhibit smaller values of *L*_int_, which also vary less with the respective lengths of the chromosomes. This behavior is consistent with the recently proposed coarsening model [8, 9, 12, 13], which suggests that *L*_int_ typically scales inversely with the CO count ⟨ *N* ⟩, unless there are few COs and *L*_int_ saturates; see Fig. 8. This suggests that species with many COs simply aborted coarsening early, leading to larger ⟨ *N* ⟩ and values of *L*_int_ that are independent of *L*, whereas species with completed coarsening exhibit only the obligate CO and 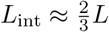. This strong connection between ⟨ *N* ⟩ and *L*_int_ also explains the observed narrow band of values of the normalized interference length 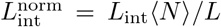 for cases where coarsening can proceed normally (e.g., in wild type and in HEI10 mutants). A similar analysis using *d*_CoC_ and *ν* does not yield a consistent picture (see Fig. 9A–D), suggesting that only *L*_int_ captures an essential property of CO interference that is nearly preserved across species.

**FIG. 8.**
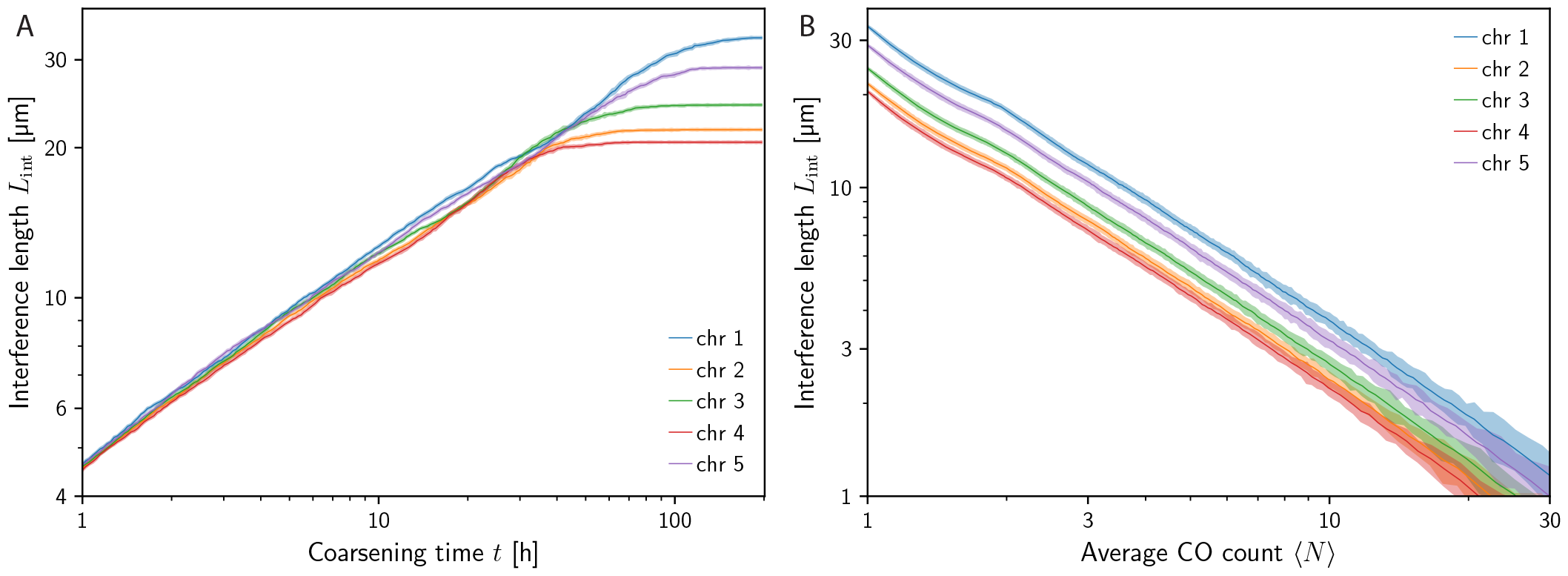
Coarsening model predicts linear growth of the interference length *L*_int_ with similar values across chromosomes and convergence to 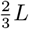 for the one CO limit. (A) Numerical data of the time evolution of *L*_int_ in the coarsening model for each individual chromosome of *A. thaliana* [12] (wild type). (B) Interference length as a function of the average CO count per chromosome ⟨*N* ⟩ for each individual chromosome.

The coarsening model explain the qualitative features unveiled by *L*_int_. Yet, the fact that it overestimates 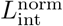 for large ⟨ *N* ⟩ indicates that additional refinements are necessary. For this, the interference length *L*_int_ will prove invaluable in quantifying the influence of model parameters and facilitate the comparison with experimental data, particular when time courses become available. At the same time, *L*_int_ can also support experimental work, particularly by allowing comparisons between species, genotypes, and chromosomes.

## AUTHOR CONTRIBUTIONS

All authors conceived the project and developed the new quantification. M.E. investigated its properties and analyzed the data. M.E. and D.Z. wrote the first draft of the manuscript, which all authors edited and approved.

## ACKNOWLEDGMENTS

We gratefully acknowledge funding from the Max Planck Society and the European Union (ERC, Emul-Sim, 101044662).

## Appendix A: Traditional quantifications

### 1. Quantification based on a Gamma distribution

To determine the shape parameter *ν* of the gamma distribution for a given set of observed data we simply apply the `python`-function `scipy.stats.gamma.fit` to the data set of all distances between adjacent COs; see Fig. 1B, Fig. 7B, and Fig. 9A.

**FIG. 9.**
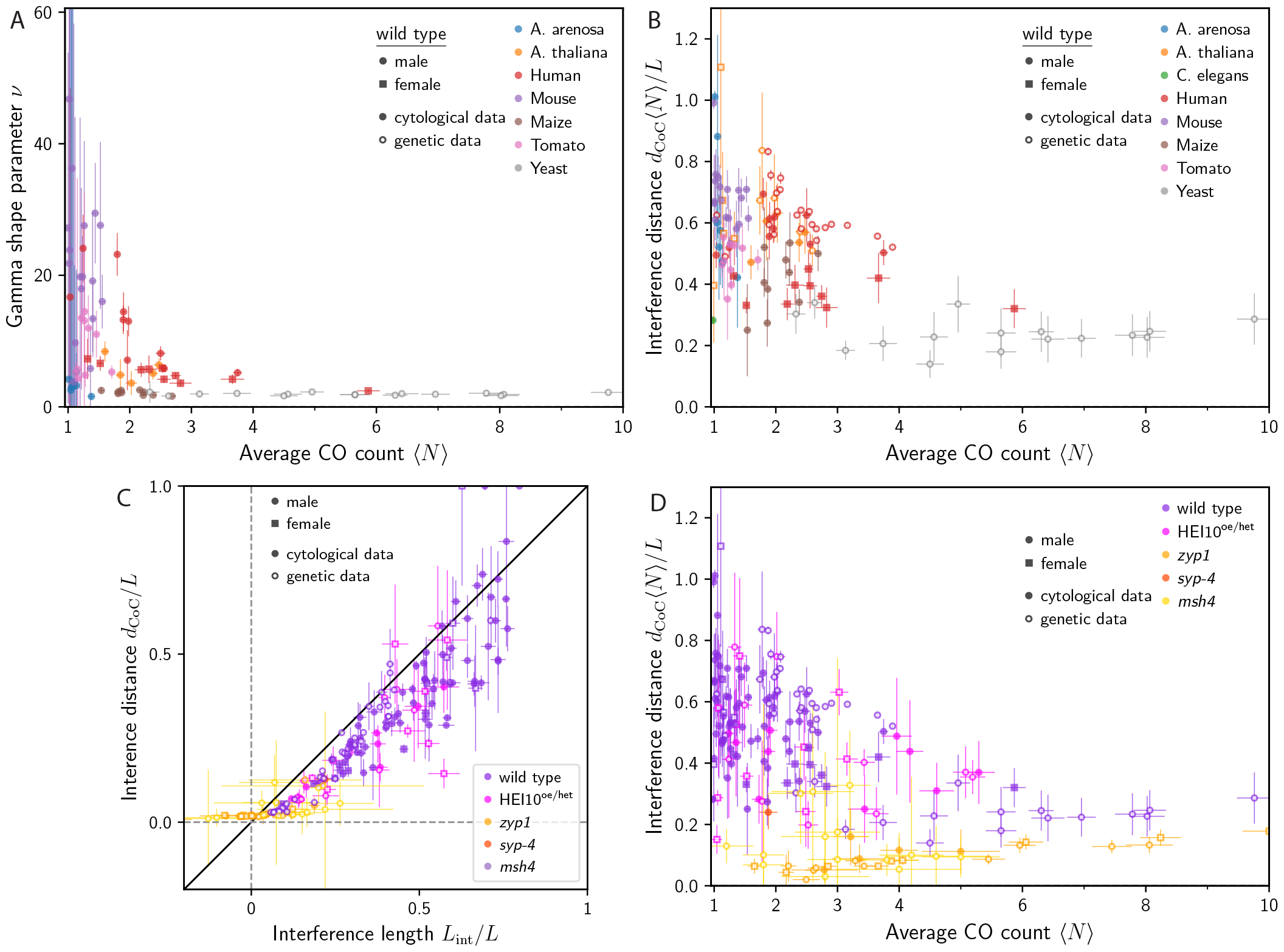
Traditional measures do not show the similarity observed for interference length. (A) Gamma shape parameter *ν* as a function of the average CO count ⟨ *N* ⟩ in logarithmic scales for wild-type data for the cytological data as in Fig. 6A and full-tetrad genetic data of budding yeast. The data of *C. elegans* is dropped because it has not enough CO pairs, as well as the genetic data of *A. thaliana* and human because the shape parameter *ν* is not invariant to random sub-sampling (cf. SI, section A 1). In particular, for small ⟨ *N* ⟩ the uncertainties are huge due to small sample sizes of observed adjacent COs. (B) Interference distance *d*_CoC_ as a function of average CO count ⟨ *N* ⟩ (using 15 intervals to compute the CoC curves) for the wild-type data for both cytological and genetic data as in Fig. 6A. (C) Correlation of interference distance *d*_CoC_*/L* and interference length *L*_int_*/L*: Comparison of interference distance *d*_CoC_*/L* and interference length *L*_int_*/L* normalized with respective chromosome length for the data points shown in Fig. 6B. (D) Interference distance *d*_CoC_ as a function of average CO count ⟨ *N* ⟩ (using 15 intervals to compute the CoC curves) for the mutant behaviour data for both cytological and genetic data as in Fig. 6B.

### 2. Coefficient of coincidence

The coefficient of coincidence (CoC) is traditionally defined by chopping up the whole length of the chromosome into intervals: For each pair of intervals *A* and *B*, the CoC is defined as the ratio of the observed frequency *r*_AB_ of double COs to the expected frequency in absence of interference [2, 3, 39]. Assuming independence of occurrence in the absence of interference, the latter is given by *r*_A_·*r*_B_, where *r*_*A*_ is the frequency of the occurrence of individual COs in interval *A*, implying [14, 38]

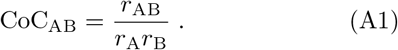

This quantity is 1 if CO positions are independent of each other (e.g., in absence of interference), whereas values smaller than 1 indicate positive interference. To summarize the data for all pairs of intervals, the CoC values are typically plotted against the distance *d* of the corresponding intervals. Since interval pairs with equal distance typically have similar CoC values, these values are averaged [38, 43]. Furthermore, the interference distance *d*_CoC_ is defined as the length *d* where CoC(*d*) first exceeds 0.5 [6, 13, 43, 44].

One shortcoming of this definition is that the number of intervals between two COs at a given distance depends on their absolute position along the chromosome and not only on their separation. In particular, the distance of a pair in two adjacent intervals can vary between zero and twice the interval length, i.e., between one interval and the next but one interval, the distances can vary from one to three times the interval length, and so on. This indicates a significant overlap of these intervals.

To avoid this problem, we refine the definition of the CoC by not dividing the entire length of the chromosome into equal intervals and then counting the COs in each interval, but rather by determining the observed and expected distribution of CO distances and then partitioning these distribution into bins. To do so, we determine the actual distances of observed CO pairs and then partition the distance distribution into bins; see the green histogram in Fig. 1A. The expected distance distribution in the absence of interference can be computed by taking all possible distances between all observed CO positions and then partition this distribution by the same set of bins; see the gray histogram in Fig. 1A. The CoC is now simply the ratio of the observed and expected distributions where the observed distribution needs to be scaled with the ratio 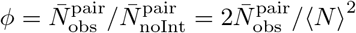 of observed to expected pairs; see definition in section II A and SI-B 1. This implementation of the CoC is used throughout our paper, i.e., in Fig. 1B, Fig. 7B and Fig. 9B–D. This implementation gives equal weight to all CO pairs and does not depend on the absolute positions of the COs along the chromosome.

## Appendix B: Properties of interference length

### 1. Number of expected pairs

We here estimate the expected number of CO pairs for the reference case of absent interference, 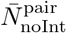, when the average CO count per bivalent is ⟨*N*⟩. In this scenario, the distribution of the number of COs per chromosome follows a Poisson distribution [39], since the COs are assumed to be independent events. Consequently, the probability of finding *m* COs in a given sample follows a Poisson distribution, 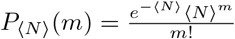. Hence,

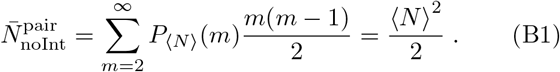

### 2. Effect of random sub-sampling

We here show that the interference length *L*_int_ is invariant to random sub-sampling of a certain fraction *φ* of the COs. First, note that the probability that a given CO *survives* after random sub-sampling is *φ*, assuming the independent probabilities for all CO. Thus, the distribution of COs along the chromosome remains unchanged, and so does the expected distribution of distances without interference, and thus also *d*_noInt_. Because the probabilities of individual COs are independent, the joint probability for any pair of COs on the same chromosome is given by *φ*^2^. Consequently, the distribution of distances of all CO pairs remains unchanged and so does the average distance *d*_obs_. This implies that the ratio *ϕ* of observed to expected CO pairs also remains unchanged,

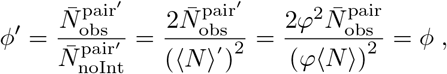

where the primed variables correspond to the values after random sub-sampling. Since *d*_noInt_, *d*_obs_, and *ϕ* are invariant under sub-sampling, *L*_int_ also remains unchanged; see Eq. (3).

If we instead considered only the distances between adjacent crossovers, the random sub-sampling would change the distribution of the distances and thus the average *d*_obs_. This justifies the choice to consider all distances *d*_*i,j*_ when determining *L*_int_.

### 3. Estimation of uncertainty

To estimate the uncertainty of the quantity *L*_int_, we determine the standard error of *L*_int_ using bootstrap sampling. To do this, we first randomly select half of the samples multiple times, calculate *L*_int_ for each collection, and determine the associated standard error of the mean. Assuming the central limit theorem, the standard error of *L*_int_ follows by scaling the bootstrapped standard error by 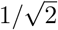 to account for the reduced sample size; see Fig. 7A.

In case of positive crossover interference, we expect the uncertainty to scale with 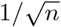. We also expect the uncertainty to be much smaller compared to absent interference since (i) the distribution of the number of crossovers per chromosome has a smaller variance (see Fig. 2A), and (ii) the distribution of the distances is also narrower; see the latter distribution in Fig. 1A. As an example, this is confirmed by the numerical simulation of the coarsening model [12] for the wild type of *A. thaliana*; see orange data in Fig. 7A.

In the following, we analytically estimate the uncertainty of the interference length *L*_int_ in the absence of interference and with the assumption of a uniform distribution of crossovers along a chromosome of length *L*. When mean CO count per chromosome is ⟨*N* ⟩ and the number of crossovers follows a Poisson distribution *P*_⟨*N*⟩_(*m*). For a Poisson distribution *P*_⟨*N*⟩_(*m*) the standard deviation of the mean of the number of crossover is given by

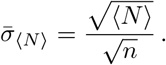

For *m* crossovers, the number of pairs is 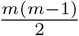 and thus the standard deviation of the number of pairs 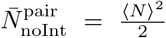 (cf. (B1)) is given by

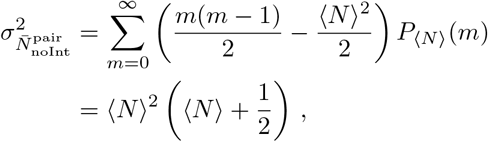

and the respective standard error of the mean by

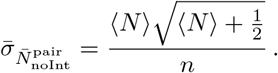

To estimate the standard error of 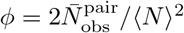 we apply error propagation including the covariance term between the number of crossover *m* and the actual number of pairs *p*_*m*_

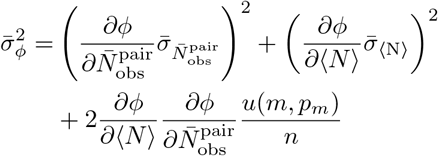

where 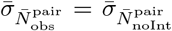 since we investigate the scenario with absent interference and covariance

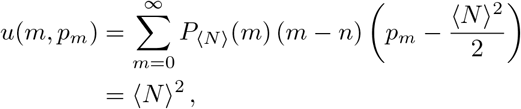

between *m* and 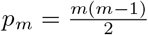, This yields

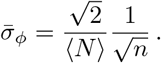

The distribution of distances of observed crossover pairs *d*_obs_ corresponds the distribution in absence of interference (cf. (B4)). Hence, we get a standard error of the mean of

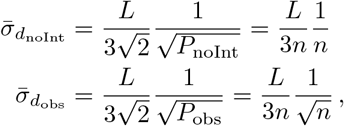

where 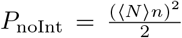 is the total number of possible shuffled pairs in the data set, and 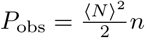 the total number of actually observed pairs. Since 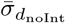 scales with 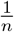 we neglect this term in the following.

Thus, in the special case of absent interference (*ϕ* = 0) and a uniform crossover distribution along the chromosome, the exact analytical estimate of the standard error of *L*_int_ (cf. (3)) is

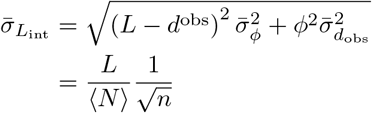

with chromosome length *L*, sample size *n*, and average CO count ⟨ *N* ⟩; see blue simulation results and black slope in Fig. 7A.

### 4. Significance testing

The paired *t*-test allows to test whether the mean of *L*_int_ of two scenarios (e.g., different mutants) across the same chromosomes in one species deviate significantly from each other. For example, we can test whether *L*_int_ exhibits sex differences, i.e., whether there is a significant difference between male and female meiosis, or compare two genotypes of the same species. In all cases, we require two scenarios (I and II), between which we want to compare *L*_int_ of the *N* ^chr^ chromosomes. For each scenario (I/II) and all chromosomes *k* = 1, …, *N* ^chr^, we calculate the interference length 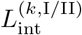 together with its standard error 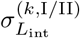 using bootstrapping. For each scenario (I/II) and all chromosomes *k* = 1, …, *N* ^chr^, we calculate the interference length 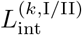 together with its standard error 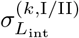 using bootstrapping. The average of the differences of this paired data set then reads

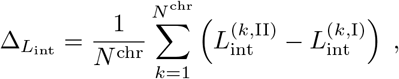

and the respective standard error of the mean is

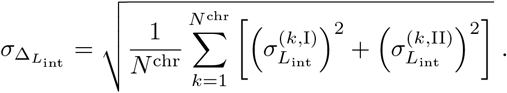

This results in the *t*-statistics,

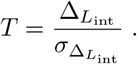

We can use this together with the null hypothesis that the interference length is identical in both scenarios to calculate the corresponding *p*-value. We use a threshold of 0.05 for rejection of the null hypothesis in our paper.

### 5. Theoretical distribution without interference

In the following, we determine the distribution of distances between all crossover pairs *d*_*i,j*_ for absent crossover interference under the assumption of uniform CO distribution along the chromosome. To do this, we distribute two points independently and uniformly along a straight line with unit length. The distribution of distances between these two points on a non-periodic line is equivalent to the distribution of distances between adjacent points if we place three points *x*_1_ *< x*_2_ *< x*3 with *x*_*i*_ ∈ [0, 1) on a line with periodic boundary conditions (a circle) since the end point corresponds to only one other random point. Without loss of generality, let *x*_1_ = 0. We then consider the cumulative probability

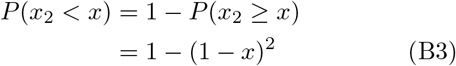

since there are two remaining points in the interval [*x*, 1). From this follows the probability distribution

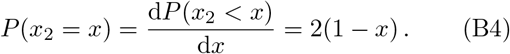

This results in an average distance of

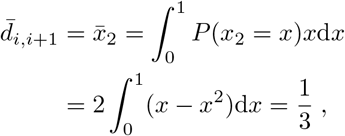

resulting in 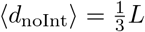 in the case of a uniform distribution. Thus, for the one-crossover limit (corresponding to *ϕ* = 0), we get 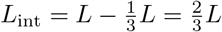.

### 6. Maximal-interference model

To investigate the maximal value of *L*_int_*/L* for given ⟨ *N* ⟩, we consider a model where exactly *N* COs are placed in regular intervals. For *N* ≥ 2, we place the two outermost COs exactly at the end of the chromosomes to obtain a normalized distance of *L/*(*N* − 1) between adjacent COs. In this scenario, we have 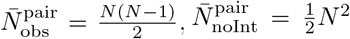, and 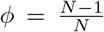. Note that the CO frequency is not uniform for *N* ≥ 2, but rather peaked. We have

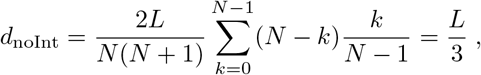

whereas

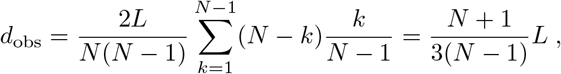

and hence

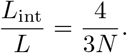

Note also that *L*_int_*/L*≤1 since *ϕ*≥ 0, *d*_obs_≤*L* and *d*_noInt_≥ 0. Hence, we predict *L*_int_*/L* = 1 for *N* = 1 (assuming a peaked CO distribution along the chromosome) in the maximal interference model. To extend the model to non-integer values of *N*, we simply interpolate to obtain a general trend,

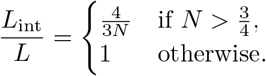

### 7. Model with regular placement

For the regular-placement model, we place exactly *N* COs in regular intervals of relative length *L/N* along the chromosome. Here, we place the first CO uniformly between 0 and *L/N*, so the overall CO frequency is uniform along the chromosome. In this scenario, the number of observed pairs is 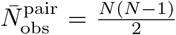 while 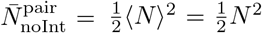, which yields 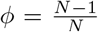. Because the CO requency is uniform along the chromosome, the average distance in case of no interference is 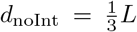; see SI-B 5. The observed average distance between pairs is given by

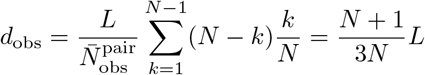

since we have *N*−*k* pairs of distance 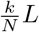. Applying Eq. (3), we get

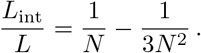

To extend the model to non-integer values of ⟨ *N* ⟩, we interpolate this relation to obtain a general trend. Note that the regular-placement model implies lower *L*_int_*/L* than would be possible for a more peaked CO distribution along the chromosome, so it does not correspond to maximal interference.

### 8. Effect o f c lass I I crossover

A challenge in interpreting crossover interference experimentally is that nearly non-interfering class II COs are indistinguishable in genetics, and invisible with MLH1 from the interfering class I COs.

In the CoC curve, this can be compensated by computing the apparent CoC [59]

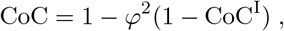

where CoC^I^ is the pure coefficient of coincidence of class I crossover as a function of distance, and 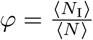 denotes the fraction of class I COs.

To determine the effect of class II COs on the interference length *L*_int_, we assume that the placement of class I and class II crossover is independent, and both classes exhibit the same distribution along the chromosome. Now, the fraction of class II COs is *φ*^II^ = 1 − *φ* = ⟨*N*_II_⟩*/*⟨*N* ⟩ (*φ, φ*^II^ ∈ [0, 1]). For simplicity, we assume that class II COs do neither show interference with other class II Cos nor with class I COs, implying there are ⟨NI⟩⟨NII⟩ mixed pairs. The number of observed pairs, 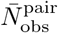, can thus be written as

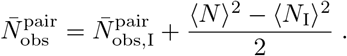

In case of absent interference, we expect 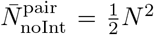 CO pairs, including 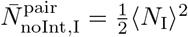 pairs of only class I COs and 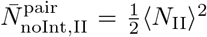 pairs of only class II COs. Defining

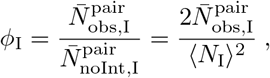

we find

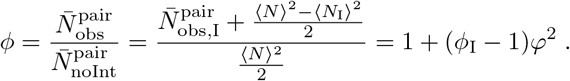

Applying the average distance of class I CO, 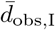,

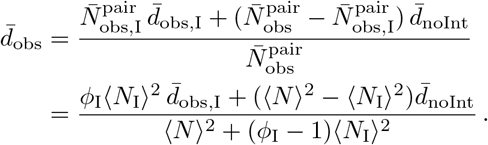

Using Eq. (3), we finally conclude

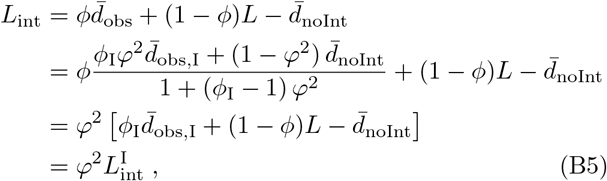

where 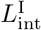 is the pure interference length measured for only class I COs. Consequently, a higher share of class II COs reduces the apparent interference length *L*_int_. However, Eq. (B5) allows to estimate the pure interference length of class I COs based on (an estimate of) the fraction of class II crossover.

This analysis assumed that the class II COs do not interfere at all. While an analysis of tomato [18] shows that class II CO indeed not interfere with each other, there might be a small but positive interference between class I and class II CO. This effect reduces the accuracy of the above estimates.

### 9. Calculating interference length for pairs of one class I and one class II CO

To quantify how class II COs interfere with the positioning of class I COs, we extend the definition of the interference length *L*_int_ to be able to handle pairs of exactly one class I CO and one class II CO for a chromosome. For that, we need the positions of both types of COs for all samples, denoted by 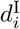 and 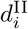. To obtain *L*_int_ from Eq. (3), we need to compute *φ, d*_obs_, and *d*_noInt_. To estimate *d*_noInt_, we compute the average of all distances between one class I COs and one class II COs over all samples. Similarly, *d*_obs_ is simply the average of only the observed distances between these mixed pairs, which also gives the number of observed pairs per sample *p*_obs_. Finally, we need estimate the ratio of observed to expected pairs, *ϕ*. The number of expected pairs is now given by

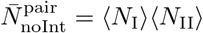 (compare SI-B 8), implying

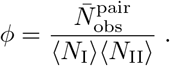

We can thus apply Eq. (3) to compute the interference length for the mixed pairs of one class I CO and one class II CO.

## Appendix C: Data

To investigate the effect of crossover interference in detail, we used empirical data sets for multiple species and genotypes. For various genotypes of *A. thaliana*, we used cytological data [9, 48] and genetic data [12, 47]. The cytological data [9] was filtered by only using the data where the column Stage was in the late phase of meiosis and the quality was 1.0. In the cytological data for the *zyp1* mutant [48], most CO position were measured twice, so we averaged those measurements. For a certain experimental line of the wild-type (wt2), the HEI10^oe^, *zyp1* and *zyp1* HEI10^oe^ mutations the chromosome number 4 was excluded from the analysis because of a potential inversion in the *Ler* line [12]. The HEI10^het^ data is not published yet, but is provided by the authors [47]. The genetic background of the mutants is not identical for the cytological and genetic data, but rather represent similar levels of HEI10 over- and underexpression; details are provided in the respective publications [9, 12, 47, 48]; compare Fig. 4.

Additionally, we used data for *A. arenosa* [45] (cytological), *C. elegans* [46] (cytological), human [50] (cytological) for both male and female, human [49] (genetic) and respective chromosome lengths from [51], maize [53] (cytological, data provided in [56]), mouse [54] (cytological, data provided in [57]), tomato [18] (cytological, data provided in [58]) and for *Saccharomyces cerevisiae*/yeast [55] (genetic data from full tetrads). For *C. elegans*, the wildtype data is given for two genotypes, with *ie29* having an HA epitope tag at the SC [46]. For most species, the data of CO positions was provided as relative position along the respective chromosome in the interval [0, 1] together with the average length of the chromosome or SC over all samples. However, for the cytological data for *A. thaliana* [12, 48] and the data for *C. elegans* [46], the CO positions were given as absolute positions [in µm] together with the SC length for each individual sample. To make these different data comparable, we also computed the relative CO positions and average SC lengths in these cases.

To determine the average CO count *N* per *bivalent* when analyzing genetic data (human [49] and *A. thaliana* [9, 12, 47]), we double the observed value to compensate for sub-sampling of COs. However, for genetic data from *Saccharomyces cerevisiae*/yeast [55] this is not necessary since the data are based on genetic analysis of full tetrads.

